# DNA methylation and hydroxymethylation quantification using vibrational spectroscopy

**DOI:** 10.64898/2026.04.02.716174

**Authors:** Rashad Fatayer, Stephen-John Sammut, Ganapathy Senthil Murugan

## Abstract

Global quantification of DNA cytosine modifications, including 5-methylcytosine (5-mC) and 5-hydroxymethylcytosine (5-hmC), is important for understanding cancer biology, though established methods require multi-step workflows and costly instrumentation. Here we show that attenuated total reflectance Fourier transform infrared (ATR-FTIR) spectroscopy combined with regression modelling enables rapid, label-free, and non-destructive quantification of both modifications from DNA samples. Using Adenomatous Polyposis Coli (*APC*) promoter DNA standards spanning 0-100% modification, we identified modification-sensitive spectral features and observed that 5-hmC produces greater spectral changes than 5-mC. A univariate peak-ratio approach yielded strong linearity for both modifications (R^2^ = 0.97), while partial least squares regression (PLSR) improved quantification accuracy to R^2^ = 0.99 (RMSE = 2.6%) for 5-hmC and R^2^ = 0.97 (RMSE = 5.7%) for 5-mC. In composite mixtures containing all three cytosine states, 5-hmC remained highly quantifiable (R^2^ = 0.97; RMSE = 5.1%), while 5-mC accuracy decreased (R^2^ = 0.90; RMSE = 9.6%), consistent with the greater spectral distinctiveness secondary to the hydroxymethyl group. Transferability was assessed using circulating tumour DNA (ctDNA), short cell-free DNA fragments shed from tumour cells into the bloodstream, comprising multiplexed reference material spanning seven genomic regions and a polydisperse fragment-length distribution (155-220 bp). After domain adaptation between synthetic and ctDNA spectra, we obtained a quantitative methylation calibration with R^2^ = 0.98 and RMSE = 5.2% under cross-validation. These results support ATR-FTIR spectroscopy as a viable platform for global cytosine modification quantification and establish proof-of-concept applicability to ctDNA analysis.

## Introduction

DNA methylation is an epigenetic modification that regulates gene expression ^1,2^ and involves the enzymatic addition of a methyl (-CH_3_) group to the 5-carbon position of cytosine (C5) residues within cytosine-guanine (CpG) dinucleotide sites, generating 5-methylcytosine (5-mC). During active DNA demethylation, 5-mC can be oxidised by ten-eleven translocation (TET) enzymes to form 5-hydroxymethylcytosine (5-hmC), which functions both as an intermediate in the demethylation pathway and as a distinct epigenetic mark associated with gene regulation ^3,4^.

Disruption of DNA methylation and hydroxymethylation is associated with several human diseases ^5–8^, including cancer ^9,10^. Across tumour types, altered 5-mC and 5-hmC levels are associated with dysregulated gene expression and disease progression ^11,12^, motivating their evaluation as biomarkers for diagnosis, prognosis, and therapeutic monitoring ^13^. In this context, circulating tumour DNA (ctDNA), short tumour-derived fragments shed into the bloodstream by cancer cells ^14^, provide a minimally invasive biomarker for tumour epigenetic assessment because it retains methylation information from the cancer of origin ^15^.

Currently, methods for quantifying DNA methylation and hydroxymethylation can be broadly divided into two categories: (i) sequence-specific approaches, which identify the genomic positions of modified cytosines, and (ii) global approaches, which measure overall 5-mC and 5-hmC abundance without positional information ^16^. Bisulfite sequencing is among one of the most widely used base-resolution methods for genome-wide methylation profiling ^17^; however, standard bisulfite conversion does not distinguish 5-mC from 5-hmC without additional chemical steps, and the harsh conversion conditions cause DNA degradation. Enzymatic methyl-sequencing (EM-seq) was developed as a less damaging solution to bisulfite approaches, using enzymatic conversion to preserve DNA integrity while enabling base-resolution methylation analysis ^18^, although it still shares the cost and infrastructure constraints associated with sequencing-based workflows ^19^. Methylation arrays provide an alternative and more cost-effective option for profiling CpG methylation across select genomic regions and are widely used in research and clinical studies, but their analysis is restricted to predefined locations and standard arrays do not resolve 5-hmC ^20^. By contrast, global methods such as liquid chromatography-tandem mass spectrometry ( LC-MS/MS) can sensitively quantify both 5-mC and 5-hmC, but typically require DNA digestion and specialist instrumentation ^21^. Indirect global immunoassays, such as enzyme-linked immunosorbent assay (ELISA) based methods, offer a more accessible and lower-cost alternative for estimating 5-mC and 5-hmC levels, but their performance can be affected by antibody specificity, cross-reactivity, and assay-to-assay variability ^22^.

Given these limitations, we investigated ATR-FTIR spectroscopy combined with machine learning as a rapid, label-free approach for global sample quantification of cytosine modifications. ATR-FTIR requires minimal sample preparation and no chemical conversion, and its low consumable requirements and straightforward instrumentation make it well suited to resource-limited laboratories seeking measurements of 5-mC and 5-hmC.

While vibrational spectroscopy has previously been shown to be sensitive to DNA methylation ^23–26^, including quantitative estimation of methylation levels in cancer cell lines ^27^, the assessment of hydroxymethylation and simultaneous determination of global 5-mC and 5-hmC in sample remain unexplored. Moreover, to our knowledge, applying such analysis to ctDNA has not previously been demonstrated using vibrational spectroscopy, despite the potential clinical value of minimally invasive epigenetic testing. Here, we present a framework for determining 5-mC and 5-hmC levels and assess its feasibility for ctDNA analysis.

To achieve this, we first characterised the ATR-FTIR spectral features of unmethylated DNA, 5-mC DNA, and 5-hmC DNA. We then evaluated two complementary quantification strategies: a univariate peak-ratio method and multivariate machine-learning models, each trained to independently estimate 5-mC and 5-hmC levels. Next, we prepared mixtures containing all three cytosine modification states (unmethylated, 5-mC, and 5-hmC) to train regression models for global sample quantification in more complex compositions. Finally, we assessed the translational feasibility in ctDNA by comparing ctDNA spectra to reference spectral profiles and applying domain adaptation, prior to constructing a calibration curve. Figure 1 summarises the workflow and presents a proof of principle.

**Fig. 1:**
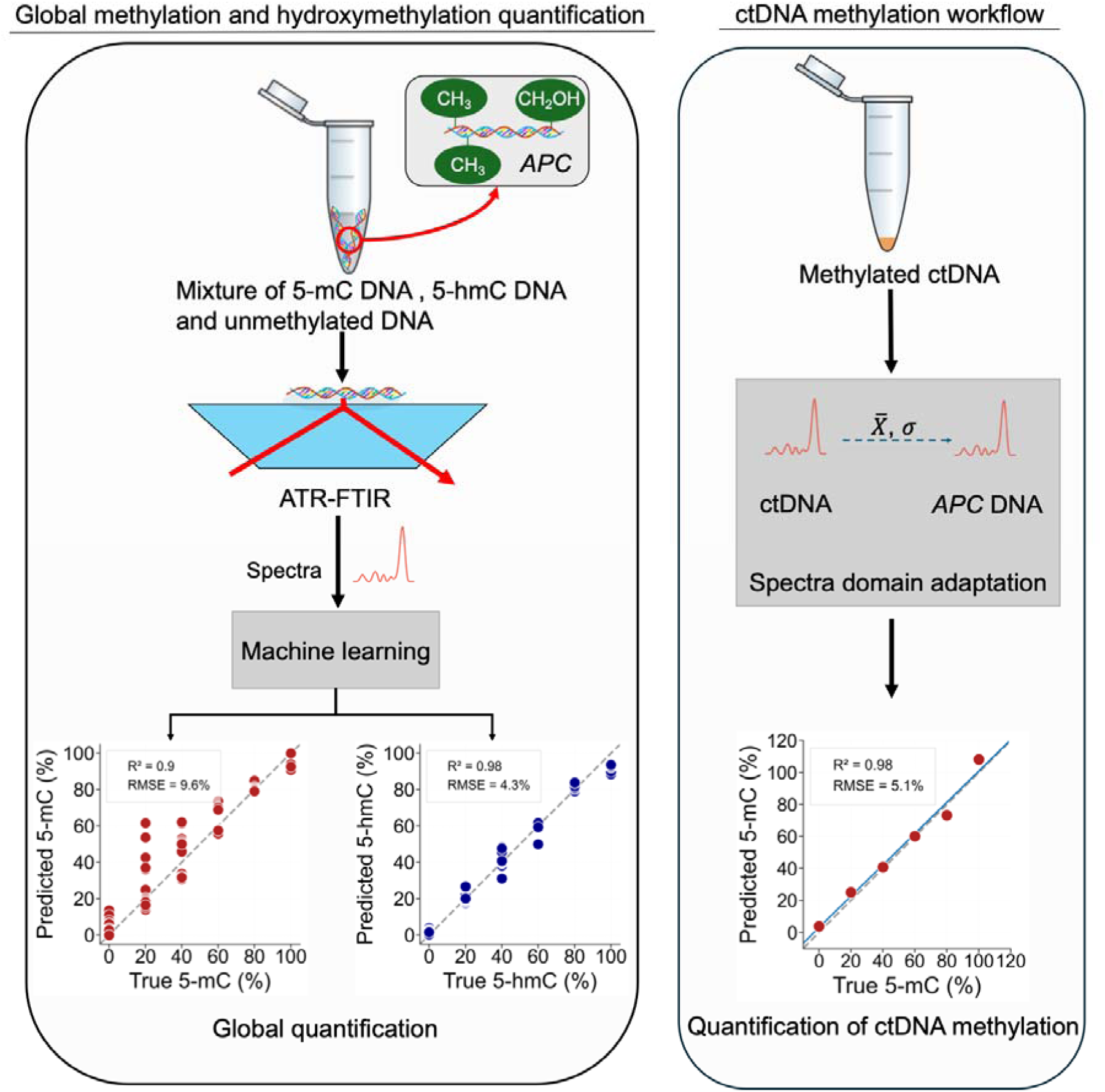
Overview of the study. Schematic workflow for global sample DNA methylation and hydroxymethylation quantification using ATR-FTIR spectroscopy and machine learning. DNA samples are measured by ATR-FTIR to generate spectral profiles, which were processed and analysed using machine learning to estimate global 5-mC and 5-hmC percentages. A parallel workflow illustrates ctDNA analysis: ctDNA spectra are acquired and domain-adapted to align with *APC* gene promoter DNA reference spectra prior to machine learning-based calibration, enabling construction of a quantitative ctDNA methylation calibration curve.

## Methods and Materials

### DNA samples

DNA standards comprising unmethylated DNA, 5-mC methylated DNA and 5-hmC methylated DNA were purchased from Active Motif (catalogue number 55008) and comprised 338 base-pairs (bp) *APC* promoter fragments generated by PCR in the presence of either deoxycytidine triphosphate (dCTP), 5-methyl-dCTP, or 5-hydroxymethyl-dCTP. Incorporation of these nucleotides during PCR produced DNA fragments containing unmethylated, fully methylated, or fully hydroxymethylated cytosine, respectively. Each double-stranded standard was provided at 50 ng/μL. The nucleotide sequence of the *APC* promoter fragment is provided in Supplementary Table 1.

Independent methylation and hydroxymethylation percentage series mixtures were prepared by mixing unmethylated *APC* promoter DNA with either fully methylated or fully hydroxymethylated DNA standards to generate samples spanning 0-100% modification in 20% increments. In each series, only one cytosine modification was varied, while total DNA concentration and volume were held constant. The 0% condition corresponded to unmethylated DNA alone, and the 100% condition to the fully modified standard. The specific mixture compositions are detailed in Supplementary Table 2.

Global cytosine modification mixtures were prepared using a simplex-based mixing strategy. Binary mixtures were first generated by combining fully methylated and fully hydroxymethylated *APC* promoter DNA standards in defined proportions to produce samples with varying global 5-mC/5-hmC ratios (60/40, 40/60, 80/20, 20/80). These samples contained only modified cytosines, with no unmethylated DNA. Ternary mixtures were subsequently prepared by introducing unmethylated *APC* promoter DNA into the binary mixtures to generate three-component compositions containing unmethylated, 5-mC, and 5-hmC DNA. Final ternary compositions included 20/20/60, 20/40/40, 40/20/40, 20/60/20, 60/20/20, and 40/40/20 (5-mC / 5-hmC / unmethylated, % of total DNA). In all cases, total DNA concentration and final volume were held constant, and proportional mixing calculations are provided in Supplementary Table 3.

### Circulating tumour DNA samples

Seraseq methylated ctDNA mutation mix reference materials (catalogue numbers 0710-3089 and 0710-3088; SeraCare) were purchased to provide high and low-methylation controls for ctDNA methylation evaluation. These reference materials comprised matched unmethylated and methylated ctDNA spanning the same seven genomic regions (*CCND2, EGFR, ETV6, FANCA, MYB, RET,* and *TFRC*). The manufacturer documentation reports low baseline methylation for the unmethylated material (∼0.44-1.1%) and high level methylation for the methylated material (>90%).

For each ctDNA material (methylated and unmethylated), a single global sample methylation value was defined by averaging the product catalogue methylation percentages reported for the seven included genomic regions. These averaged values were then used to prepare intermediate global methylation mixtures by proportionally diluting the methylated ctDNA with unmethylated ctDNA to generate methylation levels of 20%, 40%, 60%, and 80%, matching those used in the *APC* DNA experiments. The 0% condition consisted of the unmethylated ctDNA material. DNA concentration was provided at 10 ng/µL, and all mixtures were prepared at a constant final volume. The specific mixture compositions are provided in Supplementary Table 4.

### ATR-FTIR spectroscopy

ATR-FTIR spectra were acquired using an Agilent Cary 670 FTIR spectrometer equipped with a liquid-nitrogen-cooled MCT detector and a MIRacle nine-bounce diamond-coated ZnSe ATR crystal. In each measurement, 4 µL of DNA solution was deposited onto the ATR surface and air-dried at room temperature for approximately 15 min to form a uniform thin film. Spectra were collected over the range 750-6000 cm^-1^ at a spectral resolution of 4 cm^-1^, with 64 scans averaged per acquisition.

Independent replicate measurements were defined as separate depositions of the same DNA sample, in which the deposited film was removed, the ATR crystal thoroughly cleaned, and a fresh sample of the same composition of DNA solution re-deposited. This replication strategy was applied to all *APC* DNA measurements, including unmethylated, methylated, and hydroxymethylated standards, the methylation and hydroxymethylation percentage series, and the global mixture experiments, with each condition measured in triplicate. In contrast, ctDNA samples were measured using a single deposition per condition due to limited material availability. For each deposited film, nine consecutive spectra were recorded without repositioning to assess short-term measurement repeatability prior to sample removal.

### Spectra preprocessing

A coherent spectral preprocessing pipeline was applied to all *APC* DNA ATR-FTIR spectra analysed in this study. To prevent information leakage, spectra were first split into training and test sets, and all preprocessing parameters were derived exclusively from the training data and subsequently applied to the held-out test data. Spectra were truncated to 750-4000 cm^-1^ to exclude non-informative regions and vector-normalised across the retained window. A manual baseline correction was then applied to remove baseline drift artefacts using Peak Spectroscopy software. For multivariate modelling, the baseline-corrected spectra were further processed using Savitzky-Golay smoothing followed by a second-order derivative transformation ^28^ to enhance spectral resolution and improve discrimination of overlapping vibrational features.

### Statistical analysis

For comparison of relative absorbance differences between unmethylated, 5-mC, and 5-hmC *APC* DNA standards at selected spectral regions, a one-way analysis of variance (ANOVA) was performed. Statistical significance was defined as p < 0.05.

### Principal Component Analysis

Principal Component Analysis (PCA) was applied as an unsupervised dimensionality reduction technique to explore structure within the ATR-FTIR spectra and to visualise separation between cytosine modification states and methylation levels. PCA was used to identify the sources of spectral variance and to highlight regions contributing to discrimination between unmethylated, methylated, and hydroxymethylated DNA. Although five principal components were calculated, only the first two components, which captured the greatest proportion of variance, were visualised in the PCA score plots. The corresponding PC1 and PC2 loading vectors were examined to identify wavenumbers contributing most strongly to the observed separation.

To further characterise the spectral differences between unmethylated, 5-mC, and 5-hmC DNA, a Mahalanobis distance-to-centroid classifier was developed using two log-ratio spectral features derived from the cytosine modification associated bands identified in the PCA loading analysis. The features were defined as:

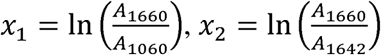

where each absorbance value was computed as the mean centred at the specified wavenumber. Classification was performed by computing the Mahalanobis distance from each spectrum to the centroid of each class, using per-class covariance matrices estimated from the training data. Each spectrum was assigned to the class whose centroid yielded the smallest distance. Classifier performance was evaluated using leave-one-group-out cross-validation (LOGO-CV), in which all scans from one independent sample composition were held out as the test set while the classifier was refit on the remaining sample compositions. The class centroids and covariance matrices were recomputed within each fold using only the training spectra, ensuring no information from the held-out composition contributed to the classifier parameters. In total, nine groups were defined across three independent sample compositions per class.

### Partial Least Squares Regression

Partial Least Squares Regression (PLSR) was used as a supervised regression method to quantify DNA methylation and hydroxymethylation levels from ATR-FTIR spectra. Data were divided into training and independent test sets using a target ratio of approximately 80:20. Splitting was performed in a replicate-aware manner, such that spectra arising from the same experimental deposition replicate, including all associated repeat scans, were assigned to only one partition. For *APC* promoter DNA samples, replicate assignments were selected and dataset sizes adjusted where necessary to preserve replicate integrity while keeping the final training:test split close to 80:20. Repeated scans from the same deposition were therefore kept together and treated as a single grouped unit during model development.

Model optimisation and latent variable (LV) selection were performed to minimise overfitting. For each model, 10-fold cross-validation was used to assess performance as a function of the number of latent variables, using the coefficient of determination (R^2^) and mean squared error (MSE). Van der Voet’s F-test was then applied sequentially between consecutive numbers of LVs to identify the lowest-complexity model whose performance was not statistically significant to models with additional components. In cases where no statistically significant differences were observed between successive models, the optimal number of LVs was selected by maximising R^2^ and minimising MSE while maintaining model parsimony. Following cross-validation, the selected number of LVs and all model parameters were fixed, and the final PLSR model was evaluated on the independent held-out test set to generate predictions.

### Domain adaptation

A domain-adapted PLSR pipeline was used to calibrate ctDNA methylation against the *APC*-derived DNA and account for spectral differences. *APC* and ctDNA spectra were interpolated onto a common 750-1800 cm^-1^ range. Model evaluation used leave-one-methylation-level-out cross-validation, where all spectra from one ctDNA methylation percentage were withheld as the validation set in each fold and the remaining methylation levels were used for training.

All preprocessing, adaptation, and modelling steps were fit using training data only within each fold and then applied to the held-out ctDNA spectra without refitting. Spectra were first smoothed using a Savitzky-Golay filter and baseline-corrected using the rubberband algorithm ^29^. To reduce DNA-type variance, PCA was fitted on the combined *APC* training spectra and ctDNA training spectra, and the PC1 associated with *APC*-ctDNA separation was removed by zeroing PC1 scores followed by reconstruction into spectral space.

After PC1 removal, mean and standard deviation alignment was applied to map ctDNA spectra onto the *APC* spectral distribution in PCA score space. A PCA model was fitted to the combined training spectra, where ctDNA scores were then centred and scaled to match the mean and standard deviation of *AP*C scores (per component) before transformation back to the full spectral domain. The resulting mean and standard deviation transform was applied to both training and held-out ctDNA spectra within each fold.

For regression, repeat ctDNA scans were averaged within each methylation level to obtain one representative spectrum per level for training, and the held-out level was averaged for validation. A PLSR model was trained to predict ctDNA 5-mC percentage from the adapted ctDNA spectra and used to predict the withheld methylation level in each fold. Cross-validated predictions across all folds were aggregated to form the ctDNA methylation calibration curve and performance was summarised using R^2^ and RMSE.

## Results and Discussion

### Characterisation of DNA cytosine modification states using ATR-FTIR spectroscopy

To investigate how cytosine modification influences the infrared spectra of DNA, three DNA standards containing unmethylated DNA, 5-mC DNA, and 5-hmC DNA were analysed using ATR-FTIR spectroscopy. All samples comprised the same 338 bp APC (adenomatous polyposis coli) gene promoter fragment at an identical concentration, ensuring that cytosine modification state was the only variable. ATR-FTIR spectra were collected over the 750-4000 cm^-1^ range and averaged for each cytosine standard (Figure 2). Prior to comparison, spectra were vector normalised across the full retained wavenumber range to reduce variation arising from differences in overall absorbance, such as those caused by sample drying or ATR contact. The 1900-2300 cm^-1^ region was excluded from analysis because of strong absorption from the diamond ATR crystal. Following normalisation, relative differences were observed across multiple spectral regions between the three standards.

**Fig. 2:**
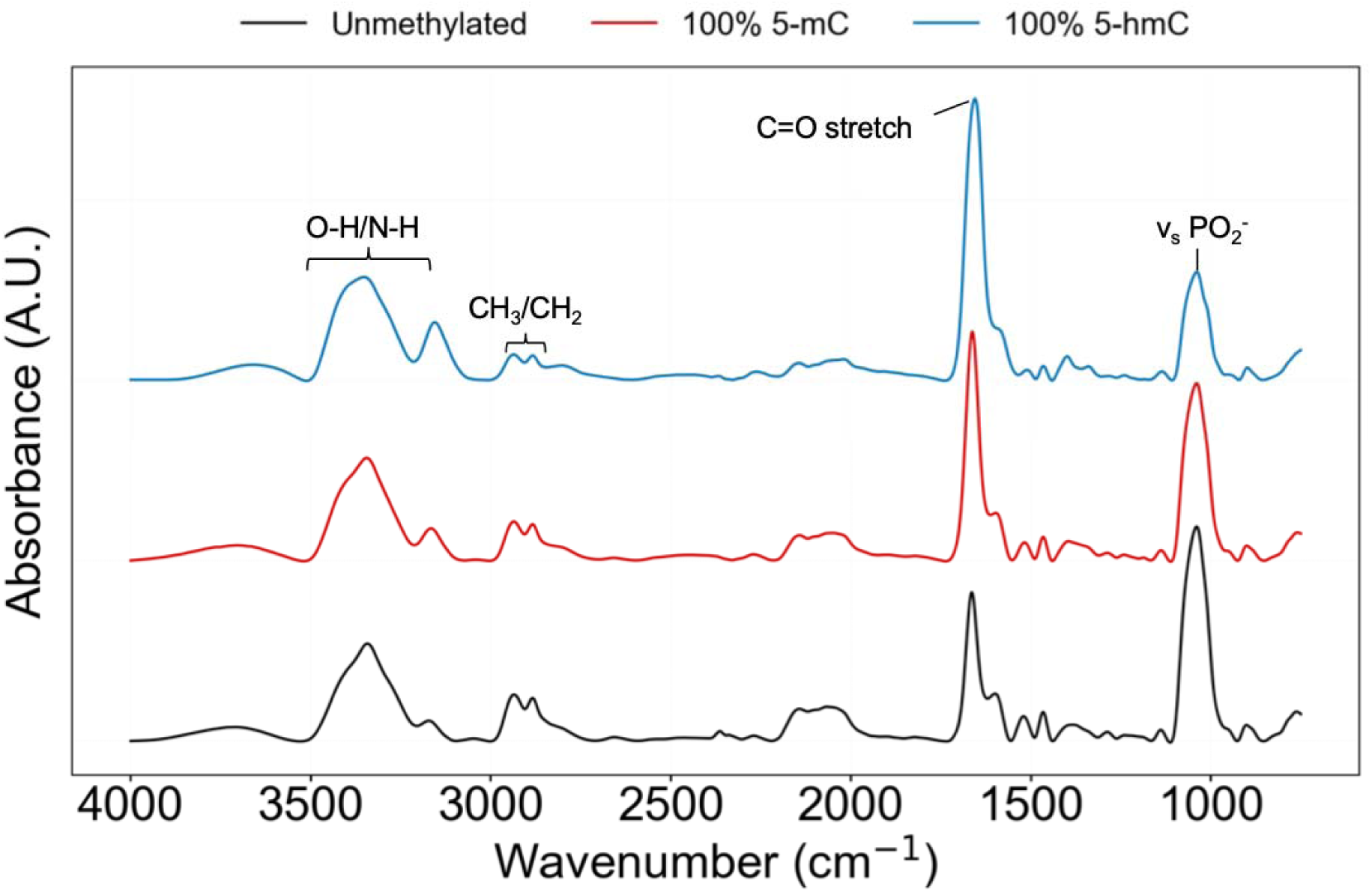
ATR-FTIR spectral analysis comparing unmethylated, hydroxymethylated, and methylated DNA samples. Full-range ATR-FTIR spectra (750-4000 cm^-1^) of unmethylated DNA (black), DNA containing 100% 5-methylcytosine (5-mC, red) and 100% 5-hydroxymethylcytosine (5-hmC, blue). Across the full spectral range, the largest modification-dependent differences are observed in four regions: (i) the phosphate backbone region (1000-1100 cm^-1^), (ii) the nucleobase C=O associated vibrations (1640-1680 cm^-1^), (iii) the C-H stretching region (2750-2900 cm^-1^), and (iv) the hydrogen bonding region (3200-3400 cm^-1^), corresponding primarily to N-H and O-H stretching vibrations.

Differences were first apparent in the phosphate backbone region (∼1000-1100 cm^-1^), where spectral features are influenced by symmetric phosphate stretching vibrations (νs PO ^-^) ^30,31^. In this region, a broad band centred at ∼1060 cm^-1^ showed both progressive decrease in absorbance intensity from unmethylated DNA to 5-mC DNA and further to 5-hmC DNA and increased band broadening in 5-hmC DNA. A one-way ANOVA confirmed a statistically significant effect of cytosine modification state in this region (*P* = 9.5 × 10^-7^). As vibrational modes in this region are sensitive to DNA conformation and hydration ^32^, these differences were most likely due to indirect effects associated with cytosine modification.

Modification-dependent changes were also observed in the ∼1640-1680 cm^-1^ region, which are associated with contributions from nucleobase carbonyl (C=O) stretching vibrations ^33,34^. The mean relative absorbance of this band increased from unmethylated DNA to 5-mC DNA and was highest in 5-hmC DNA. A one-way ANOVA again indicated a statistically significant effect of cytosine modification state in this region (*P* = 0.005). In addition, the band associated with 5-hmC DNA was broader and shifted to a lower wavenumber (1649 cm^-1^) compared with 5-mC DNA (1659 cm^-1^) and unmethylated DNA (1661 cm^-1^). Previous FTIR studies have reported wavenumber shifts within this band associated with cytosine methylation, and the trends observed here were consistent with these reports ^23,24^.

Additionally, differences associated with cytosine modification were evident in the C-H stretching region between 2750 and 2900 cm^-1^ ^30^. At approximately 2800 cm^-1^, 5-hmC DNA exhibited the highest absorbance, followed by 5-mC DNA, with unmethylated DNA producing the weakest contribution, whereas near 2875 cm^-1^ this trend was reversed, with unmethylated DNA displaying the highest absorbance. Across this region, spectral features associated with unmethylated DNA were also generally shifted toward higher wavenumbers relative to 5-mC and 5-hmC DNA. In line with prior reports, this region has been identified as sensitive to cytosine methylation ^24,35,36^, with both band absorbance and peak position varying with methylation state. The trends and peak shifts observed here therefore extend these prior observations to include 5-hmC DNA.

Finally, differences in the 3200-3400 cm^-1^ region, which included N-H and O-H stretching vibrations ^30^, further indicated modification-dependent changes. Within this region, an absorbance increase at ∼3200 cm^-1^ was observed from unmethylated to 5-hmC DNA, consistent with a direct contribution from the O-H stretching mode of the hydroxymethyl group introduced by hydroxymethylation. In addition, 5-hmC DNA exhibited a broader and more intense band centred around 3400 cm^-1^ relative to both unmethylated and 5-mC DNA, while 5-mC spectra showed little difference compared to the unmethylated control in this region. These observations for 5-hmC are likely due to the increased hydrogen bonding with surrounding water molecules, as the hydroxymethyl group is more polar than the methyl group in 5-mC.

### Validation of cytosine modification sensitive regions in DNA IR spectra using PCA

To explore whether ATR-FTIR spectroscopy could distinguish between unmethylated DNA, 5-mC DNA, and 5-hmC DNA based on their spectral profiles, and to identify the wavenumbers contributing most strongly to their differentiation, Principal Component Analysis (PCA) was applied to the preprocessed spectra. This allowed us to project the high-dimensional data into an orthogonal latent space that maximised variance capture while preserving underlying spectral features. The PCA showed clear and distinct clustering of the three cytosine modification states, indicating that each modification imparted a distinct spectral signature (Figure 3a). Principal Component 1 (PC1) accounted for 85.1% of the total variance and primarily resolved the 5-hmC samples from both unmethylated and 5-mC DNA. The second principal component (PC2), accounting for an additional 10.6% of the variance, further separated 5-mC from unmethylated DNA.

**Fig. 3:**
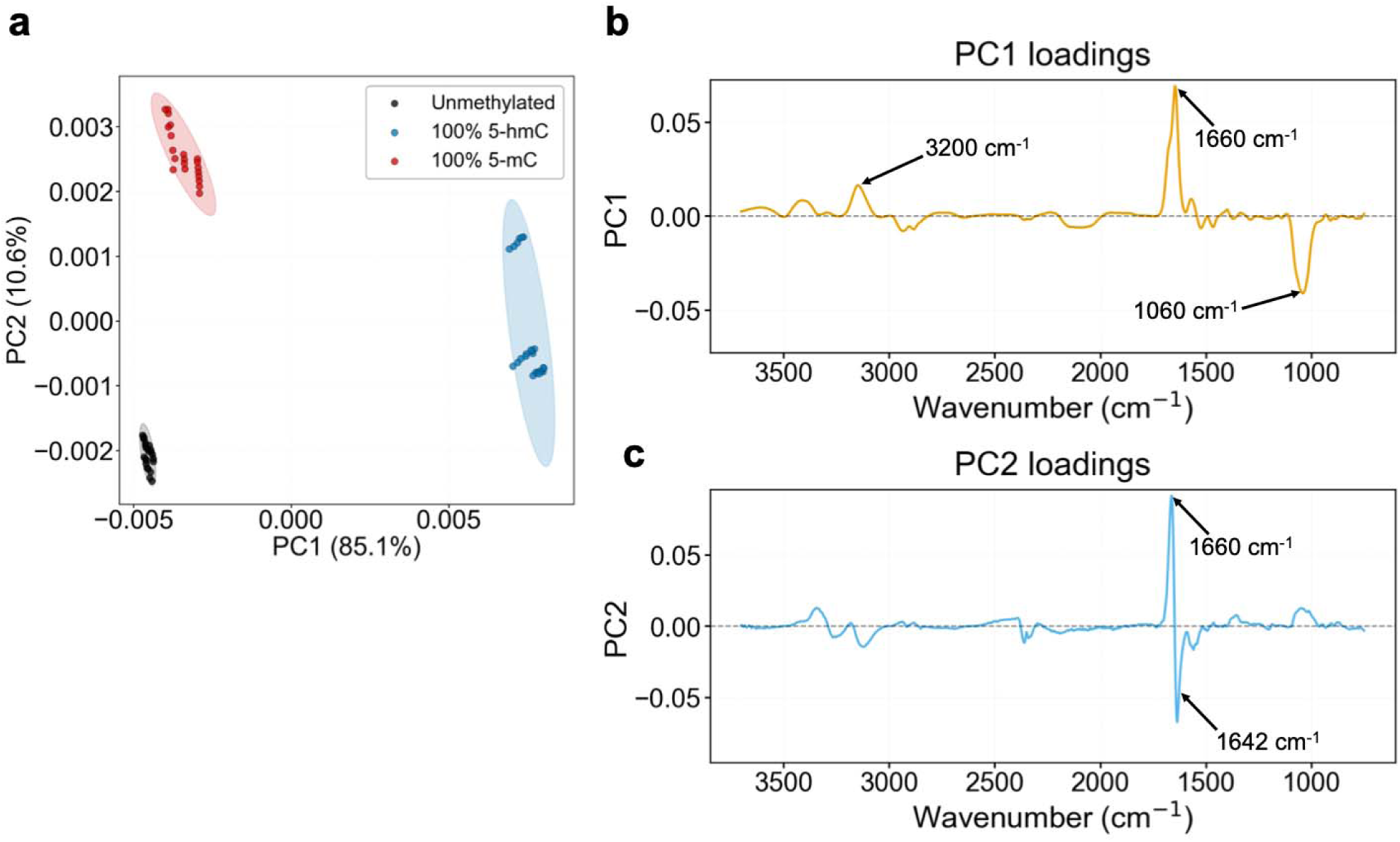
Distinguishing between DNA methylation states and regions of importance. **(a)** PCA of unmethylated (black), 100% 5-methylcytosine (5-mC, red), and 100% 5-hydroxymethylcytosine (5-hmC, blue) DNA samples, with ellipses representing the 95% confidence interval for each group. **(b)** PC1 loading plot. **(c)** PC2 loading plot. The loadings identify the spectral regions that influence the explained variance, which is related to the clustering between the groups.

We next assessed whether the observed group clustering was statistically significant using permutation testing on the PCA scores. The overall multigroup permutation test confirmed significant differentiation between the three DNA methylation states (*P* = 0.0037, 10,000 permutations). The observed test statistic, defined as the sum of squared distances between group centroids in PCA space, occurred far beyond the distribution obtained from permuted data (Supplementary Figure 1), indicating that the observed clustering pattern was unlikely to arise from random variation alone.

Additionally, the PC1 loading plot was used to identify the spectral features underlying the separation observed in the PCA, with the strongest contributions occurring between 750 and 3700 cm^-1^ (Figure 3b). Larger loading magnitudes indicate wavenumbers that contributed more strongly to this principal component. The strongest positive loading was observed near 1660 cm^-1^, corresponding to nucleobase carbonyl (C=O) vibrations, whereas the strongest negative loading was broader and centred around 1060 cm^-1^, corresponding to the phosphate backbone symmetric stretching vibration (νs PO ^-^). These features therefore made major contributions to the spectral variance underlying the clustering of the cytosine modification states. An additional strong contribution was also observed at approximately 3200 cm^-1^, although this was less pronounced than the other two features.

The PC2 loading plot (Figure 3c) showed that separation along this component was mainly driven by the nucleobase region, with a positive loading at 1660 cm^-1^ and a strong negative loading at approximately 1642 cm^-1^. Unlike the 1660 cm^-1^ feature, which was also prominent in PC1, the 1642 cm^-1^ band has been assigned to C5-methylated cytosine ^30^, indicating that PC2 captured methylation-associated spectral variation that likely contributed to the separation of 5-mC DNA from unmethylated DNA in the PCA scores plot. Minor contributions were also present in the C-H stretching region around 3000 cm^-1^ and in the phosphate and sugar backbone region between 900 and 1200 cm^-1^.

To summarise, PCA of ATR-FTIR spectra separated DNA samples containing all three cytosine modifications, with permutation testing confirming that the observed separation was statistically significant, and the PC1 and PC2 loading profiles identifying the nucleobase carbonyl and phosphate backbone bands as the primary drivers of this separation.

### Spectral classification of cytosine modification states using log-ratio features and Mahalanobis distance

To complement the unsupervised PCA separation, an interpretable classifier was developed based on two log-ratio features derived directly from the spectral bands identified in the PCA loading analysis (Methods). The first feature, *x*_1_ = *ln*(*A*_1660_/*A*_1060_), captured the relative absorbance of the nucleobase carbonyl band at 1660 cm^-1^ against the phosphate backbone band at 1060 cm^-1^, the two bands which showed strongest in PC1. The second feature, *x*_2_ = *ln*(*A*_1660_/*A*_1642_), captured the absorbance ratio between 1660 cm^-1^ and 1642 cm^-1^(C5-methylated cytosine), the two wavenumbers which contributed to separation along PC2. The log transformation was applied to stabilise variance for the purpose of classification.

Plotting the three cytosine modification states in the *x*_1_ and *x*_2_ feature space revealed complete separation of all three classes and non-overlapping clusters (Figure 4a). The class centroids showed an increase in *x*_1_ values from unmethylated (-0.26) to 5-mC (+0.37) to 5-hmC (+1.10), reflecting increasing absorbance of the nucleobase carbonyl band relative to the phosphate backbone with C5 substitution. In parallel, *x*_2_ decreased across the same order (+0.61, +0.54, +0.12), although to lesser extent between unmethylated and 5-mC DNA, when compared to 5-hmC DNA.

**Fig. 4:**
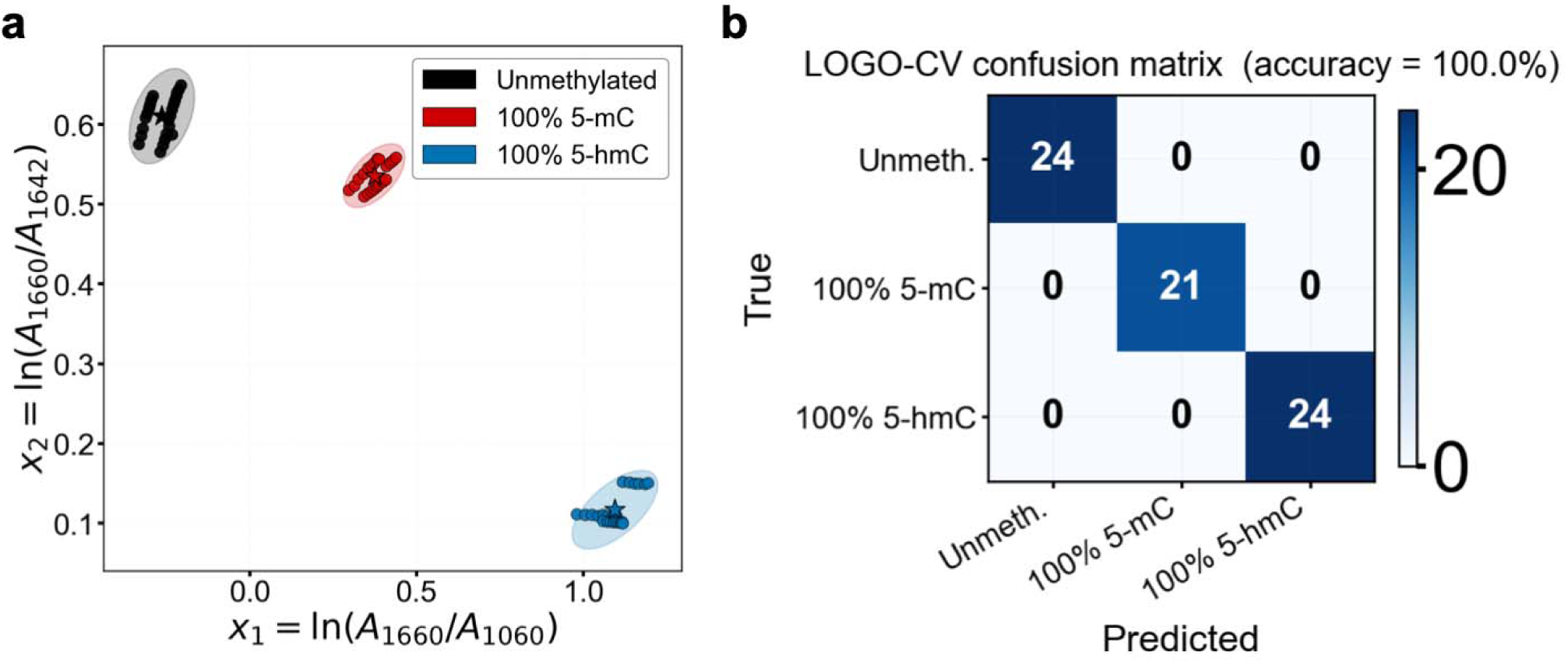
Spectral classification of unmethylated, 5-mC, and 5-hmC DNA using log-ratio features and Mahalanobis distance. **(a)** Scatter plot of two log-ratio features, *x*_1_ = *ln*(*A*_1660_/*A*_1060_) and *x*_2_ = *ln*(*A*_1660_/*A*_1642_), extracted from ATR-FTIR spectra of unmethylated (black), 100% 5-mC (red), and 100% 5-hmC (blue) DNA. Ellipses represent 95% confidence intervals for each group and stars indicate class centroids. **(b)** Confusion matrix from leave-one-group-out cross-validation (LOGO-CV), in which all scans from one independent sample composition were held out per fold. Classification was performed by assigning each spectrum to the class centroid with the smallest Mahalanobis distance, achieving 100% accuracy across all 9 folds.

Classification using Mahalanobis distance to the nearest class centroid achieved 100% accuracy under LOGO-CV, with all nine folds, where each contained a held-out independent sample composition, classified without error (Figure 4b). The large separation between class clusters relative to their within-class variance resulted in substantial margins between the assigned class distance and the nearest alternative, indicating a robust classification with no samples near the decision boundary.

### Univariate quantification of DNA hydroxymethylation and methylation levels

We next assessed a univariate quantification method based on a peak absorbance ratio of hydroxy- and methylation-sensitive spectral features. Controlled mixtures containing varying levels of 5-mC and 5-hmC were prepared as described in the Methods, in which defined proportions of fully methylated or hydroxymethylated DNA were added to unmethylated DNA to generate six modification levels (0-100%). This design enabled a systematic evaluation of how specific spectral features vary as a function of cytosine modification percentage and enabled the identification of a peak ratio suitable for quantitative analysis of both hydroxymethylation and methylation.

Progressive increases in both 5-hmC and 5-mC content produced the same inverse spectral trend, characterised by an increase in the absorbance and broadening of the nucleobase-associated band near ∼1660 cm^-1^ and a corresponding decrease in the symmetric PO_2_^-^ stretching band near ∼1060 cm^-1^ (Figure 5). For mixtures containing increasing levels of 5-hmC (0-100% in 20% increments), this trend was observed across the ATR-FTIR spectra (Figure 5a) and the equivalent spectra for mixtures containing increasing proportions of 5-mC exhibited the same relationship between these two bands (Figure 5b).

**Fig. 5:**
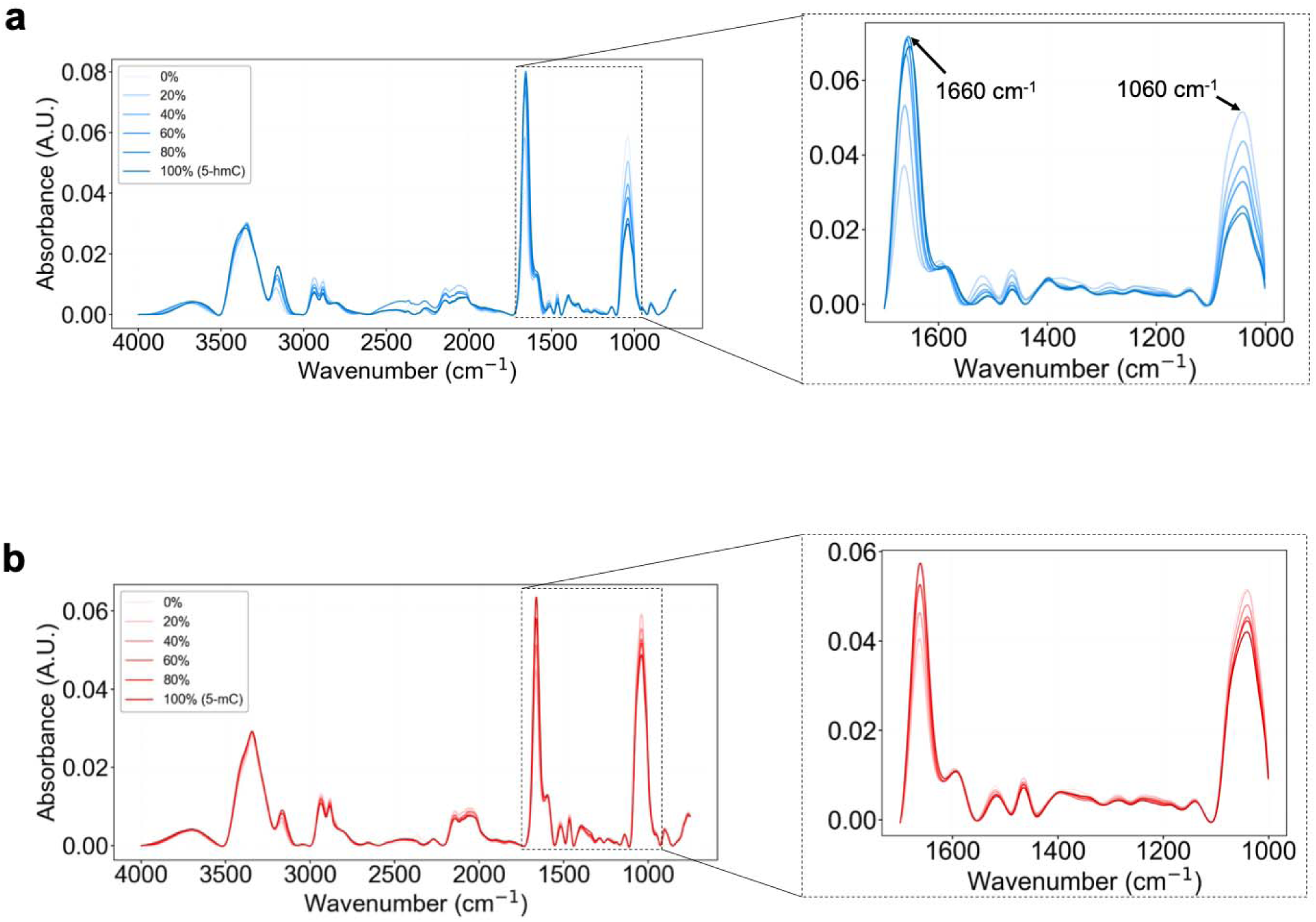
**(a)** ATR-FTIR spectra showing varying levels of 5-hmC ranging from 0% to 100% in 20% increments. **(b)** FTIR absorbance spectra showing varying levels of 5-mC ranging from 0% to 100%.

Linear regression of the 1660/1060 peak absorbance ratio against the known percentages of 5-hmC and 5-mC showed a strong relationship in both cases (R^2^ = 0.97), confirming that the relative intensities of these bands varied systematically with cytosine modification level (Figure 6a,b). The corresponding linear relationships for hydroxymethylation and methylation are given in Equations (1) and (2), respectively, where the 1660/1060 peak absorbance ratio is defined as A_1660_/A_1060_.

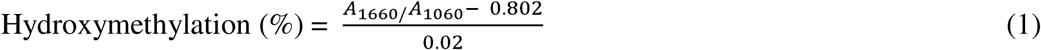

**Fig. 6:**
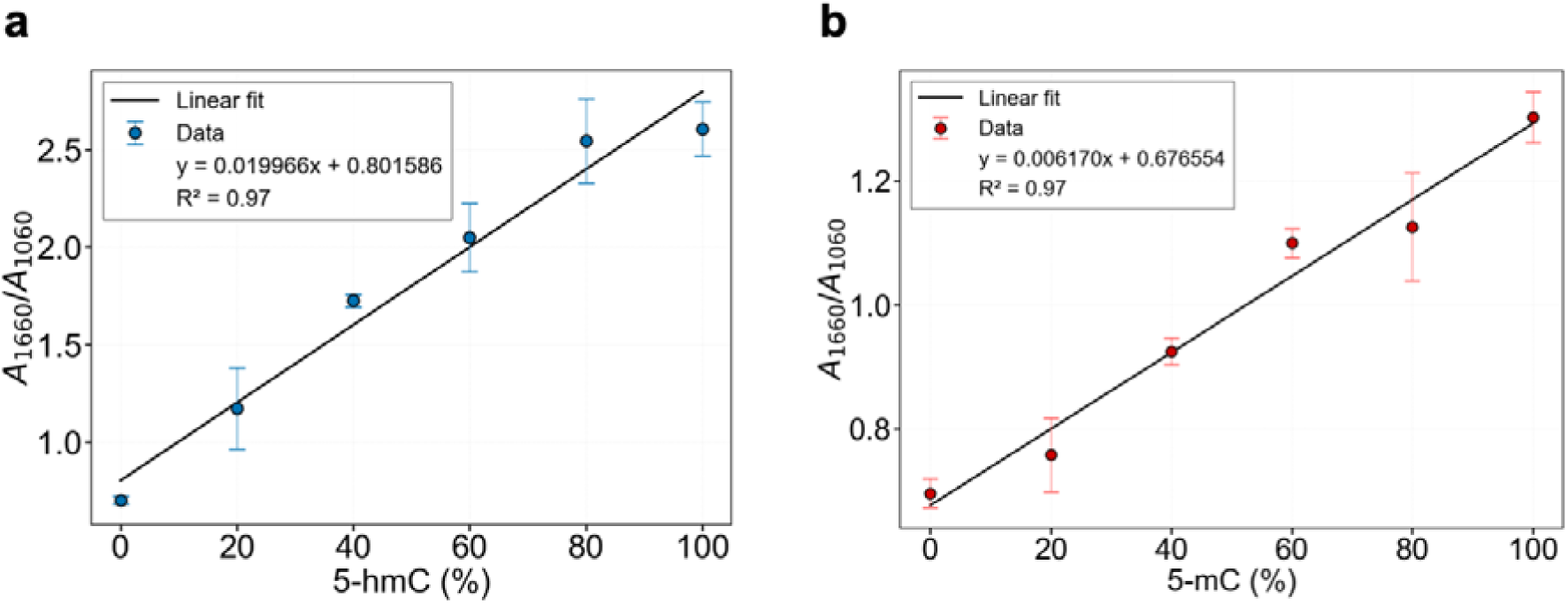
**(a)** Linear regression analysis quantifying 5-hmC levels. The plot correlates the peak absorbance ratio of 1660 cm^-1^ to 1060 cm^-1^ with the percentage of 5-hmC (R^2^ = 0.97). **(b)** Linear regression analysis quantifying 5-mC levels, plotting the same peak ratio against the percentage of 5-mC (R^2^ = 0.97). Error bars represent the standard deviation of the repeated measurements.

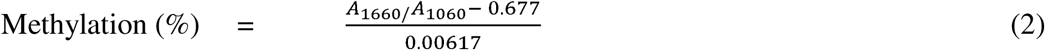

In summary, a univariate approach based on the 1660/1060 cm^-1^ peak ratio provided a simple and interpretable way to characterise variation in cytosine modification levels from ATR-FTIR spectra.

### Multivariate quantification of DNA hydroxymethylation and methylation levels

Reliance on a univariate method derived from a spectral feature can become unreliable, particularly as sample complexity increases ^37^. In contrast, multivariate machine-learning approaches incorporate information from across the spectrum, with spectral features linked through inherent collinearity. Here, we introduce Partial Least Squares Regression (PLSR), which identifies latent variables (LVs) that maximise covariance between spectral variables and cytosine modification levels, enabling quantitative prediction of methylation and hydroxymethylation percentages.

The PLSR analysis was performed over the 750-1800 cm^-1^ region, which included the most informative part of the spectrum as determined previously. Each model was developed using 163 spectra, derived from three independent spectral replicates per cytosine modification percentage. An 80/20 split was applied for model training and testing, respectively.

Two independent PLSR models were trained separately on the spectra from 5-hmC DNA and 5-mC DNA samples. Model optimisation was performed using k-fold cross-validation, with the R^2^ and the MSE to determine the optimal number of LVs (Supplementary Figure 2 and 3), followed by evaluation on a held-out test set.

The PLSR models demonstrated strong predictive performance across the full 0-100% modification range for both cytosine modifications, with the hydroxymethylation model achieving an R^2^ of 0.99 (RMSE = 2.6%) and the methylation model achieving an R^2^ of 0.97 (RMSE = 5.7%) (Figure 7a,b). The 5-hmC model consistently outperformed the 5-mC model, which may reflect the presence of more distinct spectral features in hydroxymethylated DNA arising from the hydroxyl group of the CH_2_OH moiety. In contrast, methylated DNA was spectrally more similar to unmethylated DNA, which may have reduced sensitivity for resolving intermediate methylation levels.

**Fig. 7:**
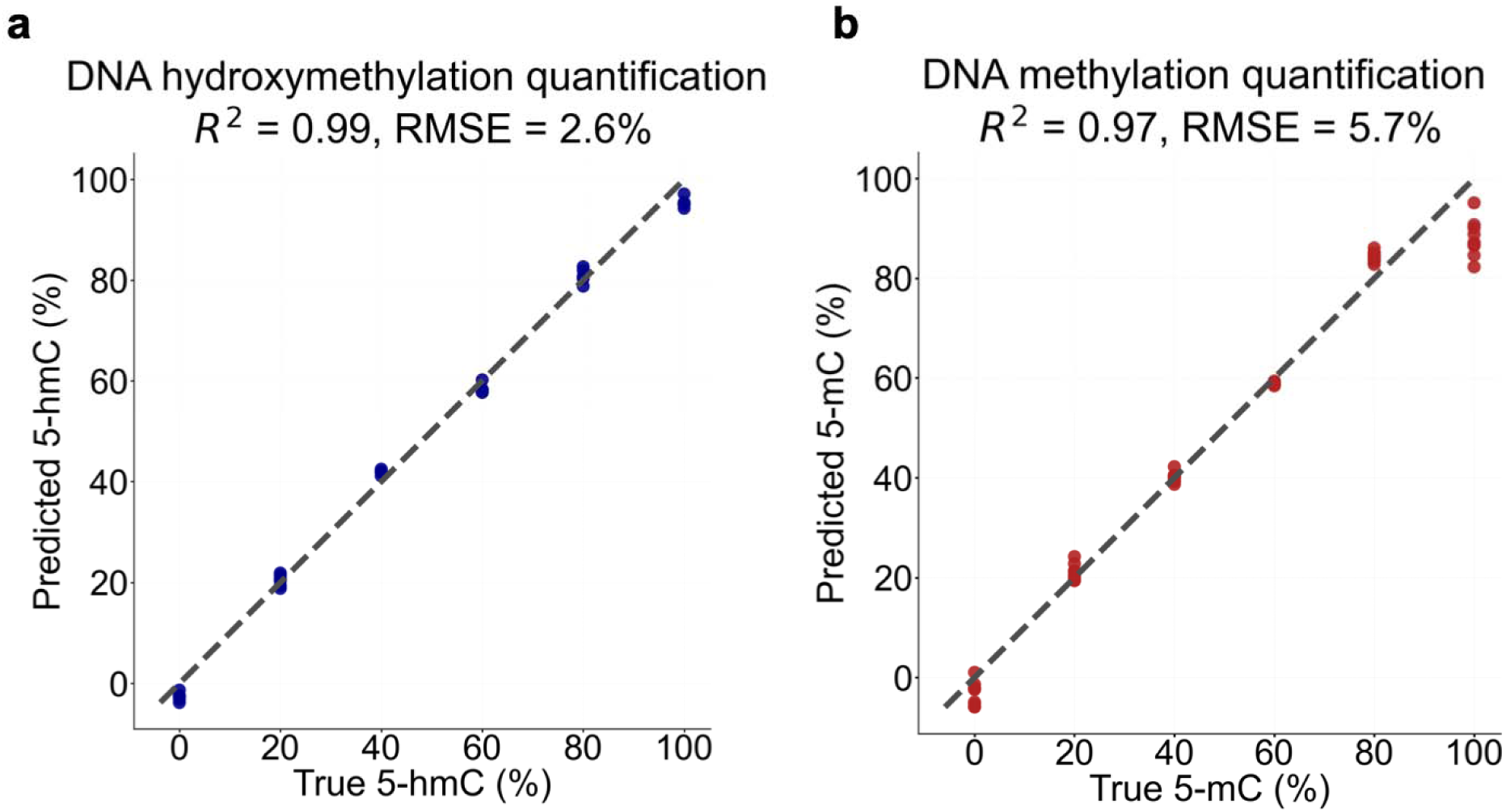
Predictions of DNA methylation levels using PLSR machine learning. **(a)** PLSR prediction plot for 5-hydroxymethylcytosine (5-hmC). **(b)** PLSR prediction plot for 5-methylcytosine (5-mC). The R^2^ and RMSE are displayed for each modification type showing predictive accuracy for 5-hmC (R^2^= 0.99, RMSE = 2.6%) and 5-mC (R^2^ = 0.97, RMSE= 5.7%).

In summary, PLSR modelling further improved predictive performance for quantifying cytosine modification levels, where 5-hmC was more accurately quantifiable than 5-mC using ATR-FTIR spectra.

### Global quantification of hydroxymethylated and methylated modification states in complex mixtures

We next evaluated a more complex mixture system in which unmethylated, 5-mC, and 5-hmC DNA were all present simultaneously within the same sample, to assess whether ATR-FTIR spectroscopy combined with machine learning could quantify global 5-mC and 5-hmC in a sample. As with earlier experiments, all mixtures were prepared from the same *APC* promoter DNA used throughout the study. By combining the three DNA standards in varying proportions, the resulting samples spanned a range of methylation and hydroxymethylation levels across the total DNA population.

Mixtures were prepared using a simplex design, incorporating ternary combinations (e.g. 20/40/40, 40/20/40, 20/60/20 for 5-mC / 5-hmC / unmethylated) and binary mixtures which contained only 5-mC and 5-hmC. Total DNA concentration and volume were held constant across all samples (Methods).

The PLSR model was trained on a combined dataset comprising spectra from the earlier single-variable series (where either 5-mC or 5-hmC was varied while the remaining fraction was unmethylated DNA) together with spectra from the new simplex mixture set. In total, the dataset comprised 433 spectra, with the same 80/20 train-test split retained.

A single PLSR model was used for the simultaneous regression of both 5-hmC and 5-mC levels within the mixture samples. Model optimisation was performed using cross validation to select the optimal number of latent variables by using the same R^2^ and MSE criteria as before, this time across the shared mixture dataset (Supplementary Figure 4 and 5).

The PLSR models maintained predictive ability under complex mixture conditions, though with clear differences between modification types. 5-hmC was quantified with high accuracy (R^2^ = 0.97, RMSE = 5.1%), indicating that the spectral features associated with the CH_2_OH group remained well resolvable even when all three modification states were present simultaneously (Figure 8a). 5-mC prediction, however, showed reduced performance (R^2^ = 0.90, RMSE = 9.6%), with scatter and mild overestimation at intermediate levels (20-60%), consistent with the more subtle spectral contribution of the methyl group becoming increasingly masked by overlapping signals from co-occurring modification states (Figure 8b).

**Fig. 8:**
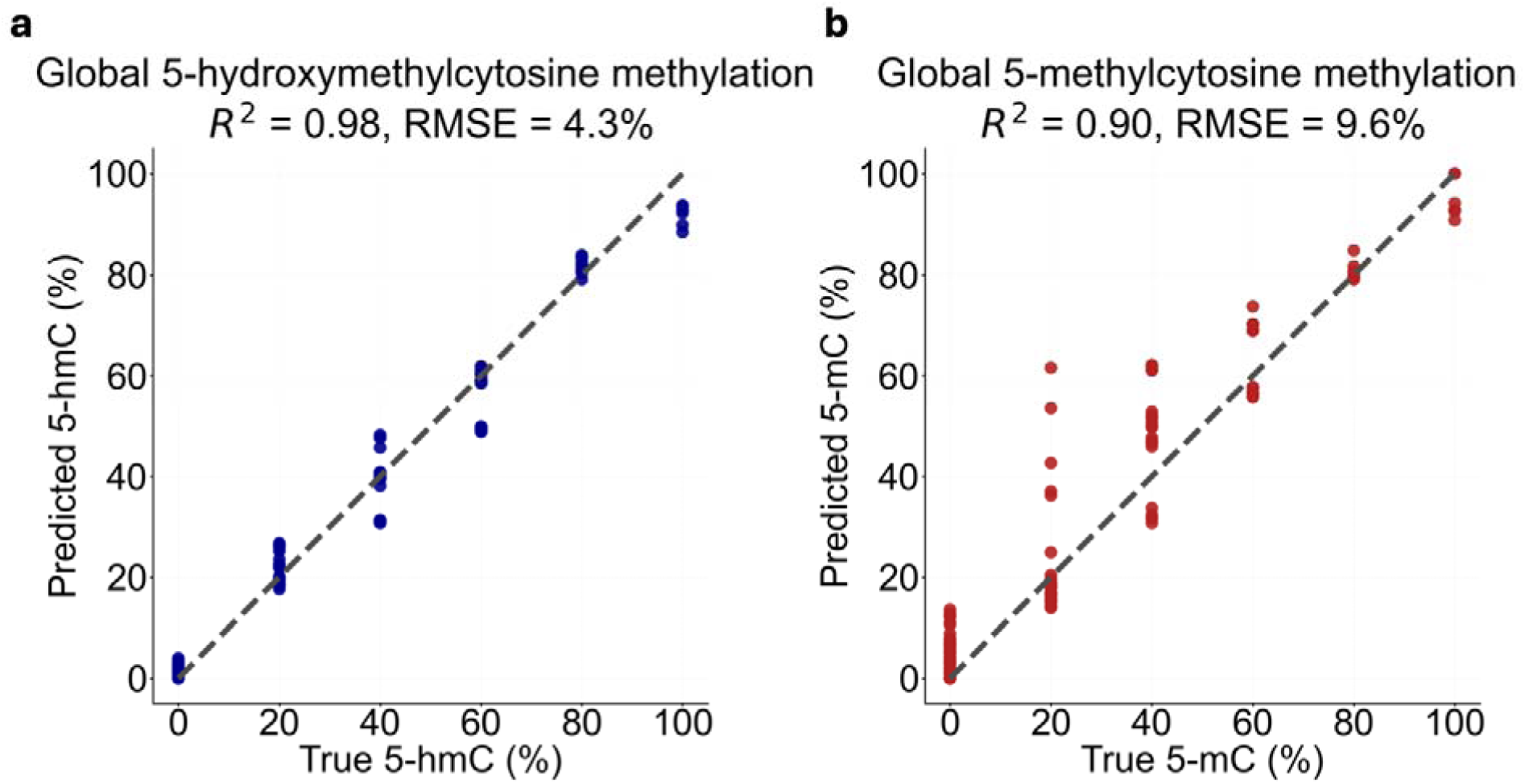
Prediction of DNA global methylation levels in complex mixtures. **(a)** PLSR quantification accuracy for 5-hmC in a mixture. Plot compares the predicted 5-hmC (%) versus the true 5-hmC (%). Data showed a strong correlation (R^2^ = 0.97) and a low error (RMSE = 5.1%), indicating high predictive accuracy for 5-hmC levels. **(b)** PLSR quantification accuracy for 5-mC. Plot compares the predicted 5-mC (%) versus the true 5-mC (%) in DNA mixtures. The model showed a lower correlation (R^2^ = 0.90) and a higher error (RMSE = 9.6%) compared to 5-hmC.

Together, these results demonstrate that ATR-FTIR spectra with PLSR can be used to quantify global cytosine modification states in mixed samples. However, quantification accuracy was affected by the spectral distinctiveness of each modification, with 5-hmC proving more resolvable than 5-mC under conditions of increasing sample complexity.

### Characterisation of circulating tumour DNA methylation levels using ATR-FTIR spectroscopy

To determine whether the methylation-associated spectral features identified translated to clinically relevant samples, a ctDNA reference standard was measured using ATR-FTIR spectroscopy. In contrast to the *APC* DNA samples, which consisted of a monodisperse fixed fragment length of 338 bp, the ctDNA reference standard comprised shorter and polydisperse continuous DNA fragments (155-220 bp) derived from a defined subset of tumour-associated genomic regions. Both methylated and unmethylated ctDNA samples were obtained, each containing seven genes (*CCND2, EGFR, ETV6, FANCA, MYB, RET,* and *TFRC*). Matched methylated and unmethylated ctDNA sharing the same genomic content were combined to produce mixtures spanning 0%, 20%, 40%, 60%, 80%, and 100% methylation, using the same mixing strategy applied in the *APC* DNA experiments (Methods). As this reference standard comprised a limited and defined subset of genomic regions, the findings presented here reflect the behaviour of this specific material and are not intended to represent ctDNA more generally without further validation.

ATR-FTIR spectra were collected for the ctDNA reference standard samples, with nine consecutive scans acquired for each of the methylation percentages, resulting in a total of 54 spectra. Due to limitations in sample availability, independent replicate measurements were not performed. The ctDNA spectra were then compared with those of the *APC* DNA samples to assess whether similar methylation-associated spectral features were retained.

Analysis of the previously identified methylation-sensitive spectral region (750-1800 cm^-1^; Figure 9a) showed that changes with ctDNA methylation level resulted predominantly in alterations in band shape, particularly band broadening, rather than a monotonic change in relative absorbance. Band broadening was also observed in the *APC* DNA series; however, in that case it progressed more systematically with increasing methylation percentage. The less uniform behaviour seen in the ctDNA reference standard likely reflected factors specific to this sample matrix, including lower analyte concentration, continuously distributed fragment lengths, and increased sequence diversity relative to the APC DNA standard, all of which would be expected to influence the relative prominence and shape of vibrational bands.

**Fig 9:**
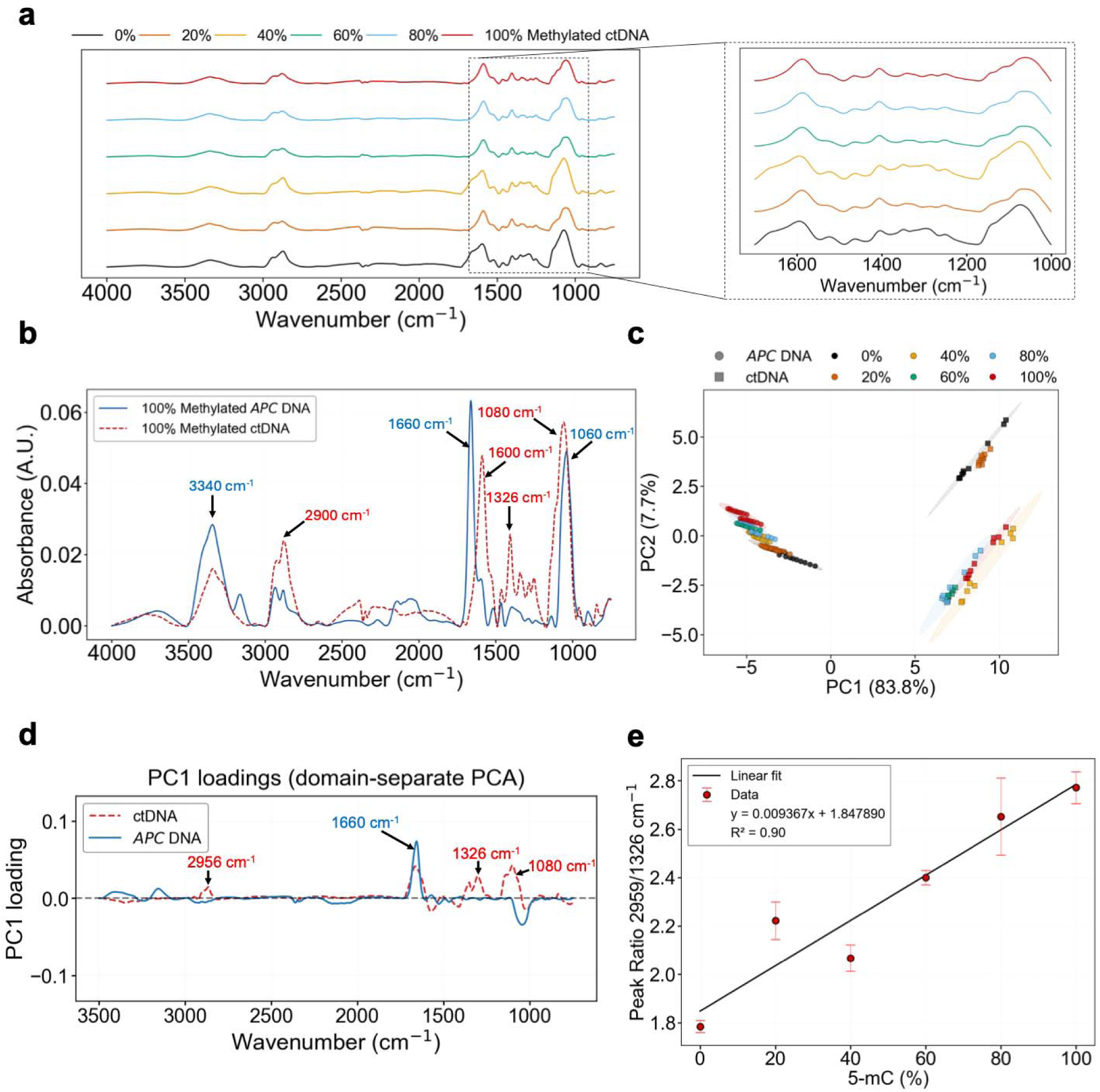
Comparative ATR-FTIR analysis of ctDNA methylation and identification of a ctDNA-specific univariate ratio. **(a)** ATR-FTIR spectra of ctDNA mixtures spanning 0-100% methylation, with the fingerprint region highlighted. (**b)** Overlay of averaged methylated ctDNA and *APC* promoter DNA spectra, showing conserved features alongside shifts in key nucleobase and phosphate-associated bands. **(c)** PCA score plot of combined *APC* DNA and ctDNA spectra, demonstrating a domain shift between DNA types along PC1 and altered methylation-related clustering behaviour in ctDNA. (**d)** PC1 loading vectors from domain-separate PCA, revealing differences in spectral features associated with changes in methylation percentage. **(e)** Calibration of the ctDNA specific 2956/1326 cm^-1^ peak ratio versus 5-mC percentage, showing a linear relationship (R^2^ = 0.90).

Direct comparison of the 100% methylated ctDNA reference standard with 100% methylated *APC* DNA (Figure 9b), together with the corresponding comparison between unmethylated ctDNA and unmethylated *APC* DNA (Supplementary Figure 6), showed that the same DNA-associated bands were present in both materials, but with clear differences in their relative prominence. In the high-wavenumber region, the broad O-H/N-H stretching contributions were more prominent in *APC* DNA (around ∼3340 cm^-1^), with the feature near ∼3100 cm^-1^ also more evident in *AP*C DNA, whereas the ctDNA reference standard showed stronger C-H stretching bands in the ∼2800-3000 cm-1 region (around ∼2956 cm^-1^ and ∼2876 cm^-1^). Within the methylation-associated region, *APC* DNA showed a stronger contribution near ∼1660 cm^-1^, while the ctDNA reference standard had a stronger contribution near ∼1600 cm^-1^. In the phosphate-associated region, both spectra contained a strong band in the ∼1100-1000 cm^-1^ range, but the dominant contribution was more prominent near ∼1080 cm^-1^ in the ctDNA reference standard compared with ∼1060 cm^-1^ in *APC* DNA. Additional differences were also apparent in the fingerprint region, where *APC* DNA showed a band near ∼1466 cm^-1^, while the ctDNA reference standard had comparatively stronger features around ∼1406 cm^-1^, ∼1326 cm^-1^, and ∼1252 cm^-1^.

PCA of the combined *APC* DNA and ctDNA reference standard spectra confirmed a domain shift between the two sample types (Figure 9c). *APC* DNA and ctDNA reference standard samples clustered on opposite sides of the score plot, with the separation occurring mostly along PC1 (83.8% variance), indicating that differences between sample type outweigh the methylation-related variance. For *APC* DNA, methylation percentage was primarily described along PC2, where samples showed well-ordered separation consistent with increasing methylation percentage across the full 0-100% range. In contrast, ctDNA reference standard samples did not show the same ordered progression with methylation percentage. Lower methylation levels (0% and 20%) clustered closely together, while samples spanning 40-100% methylation also clustered, with their separation occurring primarily along PC2. Additionally, ctDNA reference standard samples showed substantially greater within-class dispersion, indicating increased spectral variability at each methylation level within repeat measurements.

Evaluating the PC1 loading vectors obtained from PCA performed separately on the two datasets revealed differences in the spectral features associated with changes in methylation percentage between *APC* DNA and the ctDNA reference standard (Figure 9d). In the ctDNA reference standard, positive loading contributions were observed from bands near ∼2956 cm^-1^ and ∼1326 cm^-1^, which were not present in the *APC* DNA PC1 loadings. The phosphate-associated band near ∼1080 cm^-1^ also contributed positively and strongly to the ctDNA reference standard loadings, whereas this region contributed negatively in *APC* DNA. Notably, the nucleobase carbonyl band near ∼1660 cm^-1^ contributed in both datasets, indicating this region as a shared methylation-associated spectral feature across the two DNA standards.

Based on the ctDNA reference standard PC1 loadings, univariate peak absorbance ratios were evaluated using spectral features that varied most strongly with methylation percentage in this sample. Among these, the bands near ∼2956 cm^-1^ and ∼1326 cm^-1^ were selected to define a peak ratio for this reference standard. The ∼2956 cm^-1^ band was associated with C-H stretching vibrations of the methyl group of 5-mC, while the ∼1326 cm^-1^ region reflected nucleobase vibrational modes that were sensitive to methylation-associated structural changes ^23^. The 2956/1326 cm^-1^ peak ratio increased progressively with increasing 5-mC content across the ctDNA reference standard mixtures, producing a strong linear relationship (R^2^ = 0.90) and indicating that this ratio tracked methylation-dependent spectral variation within this standard (Figure 9e). Although most points followed the overall trend, the 20% 5-mC sample deviated from the fitted line, consistent with its behaviour in the PCA plot.

### Quantitative calibration of ctDNA methylation using domain-adapted PLSR

Directly utilising the methylation models developed using *APC* DNA on ctDNA would be challenging due to the observed domain shift. As shown in Figure 9c, this difference was captured predominantly along PC1 in the latent space, which separated *APC* DNA and ctDNA and reflected mostly DNA-type variance. To account for this, we developed a quantitative calibration model for ctDNA, using a pipeline integrating domain adaptation and PLSR (Figure 10a).

**Fig. 10:**
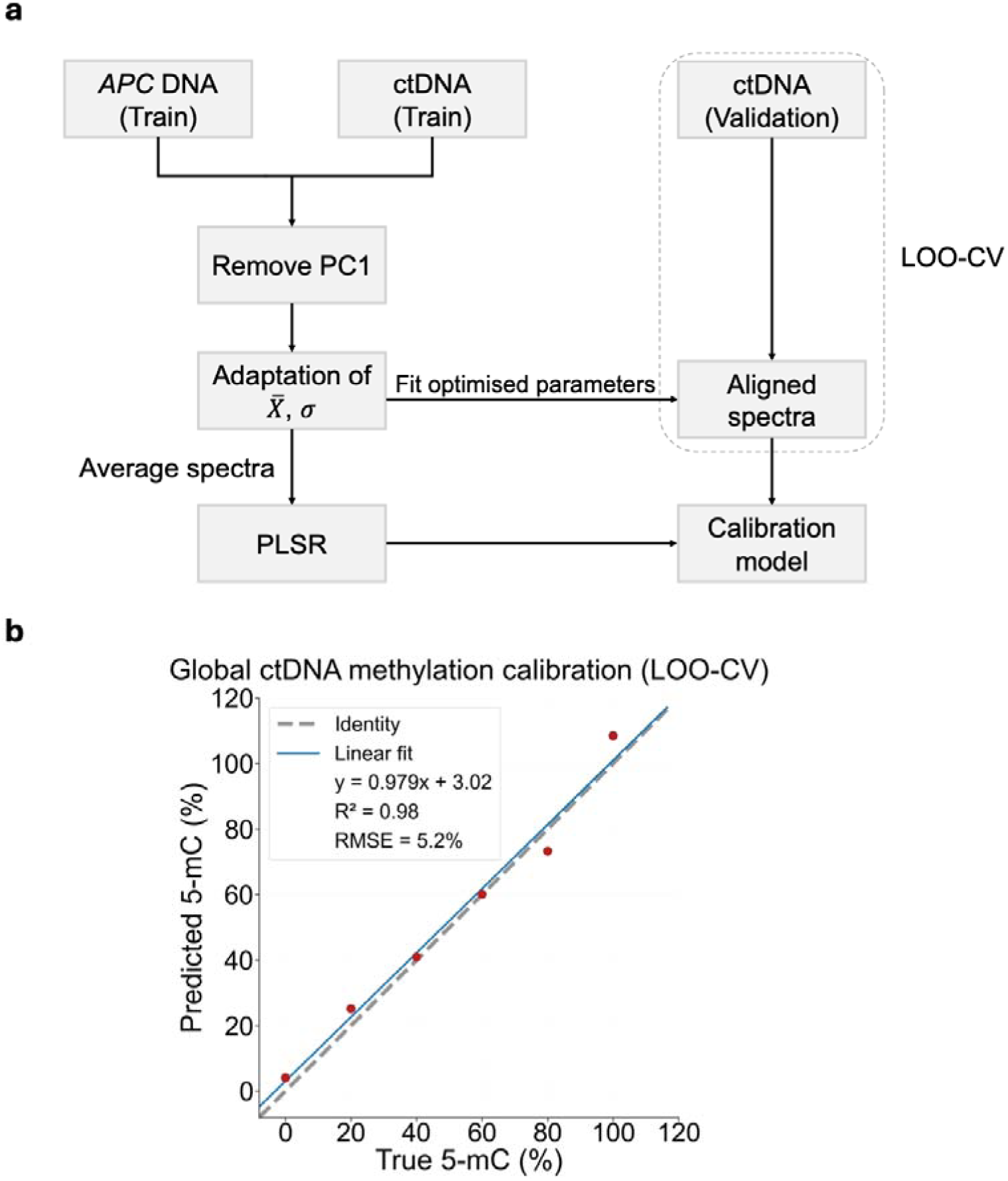
Machine learning analysis and quantification of circulating tumour DNA methylation levels. **(a)** Schematic of the domain-adapted PLSR pipeline. *APC* promoter DNA and ctDNA training spectra are used to fit optimised adaptation parameters (X□, σ), aligning ctDNA spectra to the *APC* spectral domain. Averaged adapted spectra are passed to a PLSR model to generate a calibration model, which is evaluated on held-out ctDNA validation spectra using leave-one-methylation-level-out cross-validation (LOO-CV). **(b)** Cross-validated calibration curve for global sample ctDNA methylation quantification. Predicted 5-mC (%) is plotted against true 5-mC (%) across the 0-100% range. The dashed line represents the identity relationship, and the solid line shows the linear fit to the cross-validated predictions, yielding R^2^ = 0.98 and RMSE = 5.2%, demonstrating good quantitative calibration of ctDNA methylation levels following domain adaptation.

For model evaluation, a leave one methylation level out cross-validation methodology was used. In each cross-validation fold, one ctDNA methylation level was withheld entirely as an independent validation set, while spectra from the remaining methylation levels were used for training*. APC* DNA spectra and ctDNA spectra corresponding to the training methylation levels were used to fit a PCA transformation. The domain-separating PC1 component, previously shown to capture DNA-type variance, was removed (Methods). A mean (X□) and standard deviation (σ) adaptation strategy was then applied to align the ctDNA spectra to the *APC* DNA spectra. All PCA, alignment, and modelling parameters were estimated exclusively from the training data and subsequently applied to the held-out ctDNA spectra without refitting.

Within each cross-validation fold, repeat scans of ctDNA spectra at each training methylation level were averaged to produce a single representative spectrum per concentration, which was used to train the PLSR calibration model. This procedure was repeated for each methylation level, such that in every fold the model was trained on averaged spectra from all but one methylation percentage and evaluated by predicting the averaged spectrum of the held-out methylation percentage to generate a ctDNA 5-mC calibration curve.

The cross-validated calibration model showed strong agreement between predicted and true 5-mC percentages across the full 0-100% range, producing an R^2^ of 0.98 and RMSE of 5.2% (Figure 10b), demonstrating that domain adaptation enabled a quantitative calibration prediction of ctDNA methylation levels from ATR-FTIR spectra.

## Conclusion

Here we demonstrated, for the first time, that ATR-FTIR spectroscopy can quantify both 5-mC and 5-hmC in a single, label-free measurement without chemical conversion. While the importance of these modifications in cancer is well established, existing quantification platforms require multi-step workflows, or costly instrumentation that limits their use with low-input or clinically derived material. The present work addressed this gap directly, providing a rapid and non-destructive framework applicable to both synthetic DNA standards and ctDNA reference material as a proof of principle.

Using *APC* promoter DNA fragments, cytosine modification-associated differences were observed across multiple spectral regions, most prominently near the nucleobase carbonyl band (∼1660 cm^-1^) and phosphate PO_2_ ^-^ stretching band (∼1060 cm^-1^), with 5-hmC producing the largest spectral changes. While earlier vibrational spectroscopy studies reported sensitivity to 5-mC and demonstrated some capacity for its quantification, our results showed that these same spectral regions additionally captured 5-hmC, enabling joint analysis within a single acquisition ^24,26,36^. Both a univariate peak-ratio approach and PLSR modelling successfully quantified individually varying modification levels, with the hydroxymethylation model consistently outperforming the methylation model across both strategies.

To assess true global sample quantification, binary and ternary simplex mixtures containing all three cytosine states simultaneously were evaluated using a single multi-output PLSR model. Hydroxymethylation prediction remained strong while methylation accuracy decreased, reflecting the chemical distinctiveness of each modification: the hydroxymethyl group introduced a vibrational signature that remained resolvable in complex mixtures, whereas the methyl group produced more subtle spectral changes that became increasingly masked when multiple modification states co-existed. Raman spectroscopy has similarly been shown to detect 5-hmC more readily than 5-mC ^38^, in line with the hydroxymethyl group presenting more distinct spectroscopic features across vibrational spectroscopy modalities.

We then explored clinical transferability using a multiplexed ctDNA reference material spanning seven genomic sequences and a polydisperse fragment-length distribution (155-220 bp). ctDNA spectra exhibited the same overall vibrational bands as *APC* DNA but with systematic absorbance differences, producing a measurable domain shift confirmed by PCA. Despite this, explicit domain adaptation combined with PLSR yielded a quantitative ctDNA methylation calibration with R^2^ = 0.98 and RMSE = 5.2% under cross-validation, demonstrating proof-of-concept transferability to a clinically relevant sample.

Taken together, these results demonstrated that ATR-FTIR spectroscopy with machine learning enabled accurate quantification of 5-mC and 5-hmC in controlled DNA samples and could be extended to a ctDNA reference standard when the domain shift was explicitly modelled. Future work should expand training to incorporate greater sequence and fragment-length diversity, develop ctDNA-specific models trained solely on ctDNA spectra to reduce reliance on domain transfer, and explore multimodal fusion of ATR-FTIR and Raman spectroscopy, which may improve sensitivity to 5-mC in complex mixtures by combining the complementary vibrational information of both techniques.

## Acknowledgements

This work was supported by the UK Engineering and Physical Sciences Research Council (EPSRC grant EP/S03109X/1). R.F thanks EPSRC DTP PhD studentship. We thank Breast Cancer Now for funding this work as part of Programme Funding to the Breast Cancer Now Toby Robins Research Centre. S-J.S. is a Lister Institute Prize Fellow.

## Author information

G.S.M. and S-J.S. directed the research. R.F. designed and performed the DNA methylation experiments, as well as the data analysis. G.S.M. provided funding and overall project supervision. All authors contributed to manuscript writing and approved the final version.

## Data Availability

All data supporting this study are available from the University of Southampton repository at:

## Ethics declarations

## Competing interests

The authors G.S.M, S-J.S and R.F are inventors of a patent application related to methods for analysing DNA samples (UK priority patent application GB2414133.5, filed 26 September 2024).

## Additional Information

Supplementary Information is available for this paper.

