## Supplementary Information for "DNA methylation and hydroxymethylation quantification using vibrational spectroscopy"

**Supplementary Table Legends:**

**Supplementary Table 1:** The 338 base-pair sequence of the *APC* gene used to generate the double stranded DNA standards used in the study.

**Supplementary Table 2**: Mixture compositions for the independent 5-mC and 5-hmC *APC* DNA percentage series (0-100% in 20% increments), prepared by mixing unmethylated DNA with fully methylated or fully hydroxymethylated standards at constant total DNA concentration and volume.

**Supplementary Table 3**: Composition of simplex-based global cytosine modification mixtures, including binary 5-mC/5-hmC standards (no unmethylated DNA) and ternary mixtures formed by adding unmethylated *APC* DNA; all mixtures were prepared at constant total DNA concentration and final volume.

**Supplementary Table 4:** Composition of the ctDNA methylation mixtures prepared by proportionally mixing methylated and unmethylated reference materials (*CCND2, EGFR, ETV6, FANCA, MYB, RET, TFRC*) to generate target global methylation levels (0-100, in 20% increments) at constant DNA concentration (10 ng/µL) and final volume, using manufacturer-specified regional methylation values averaged across loci to define global reference levels.

**Supplementary Figures**

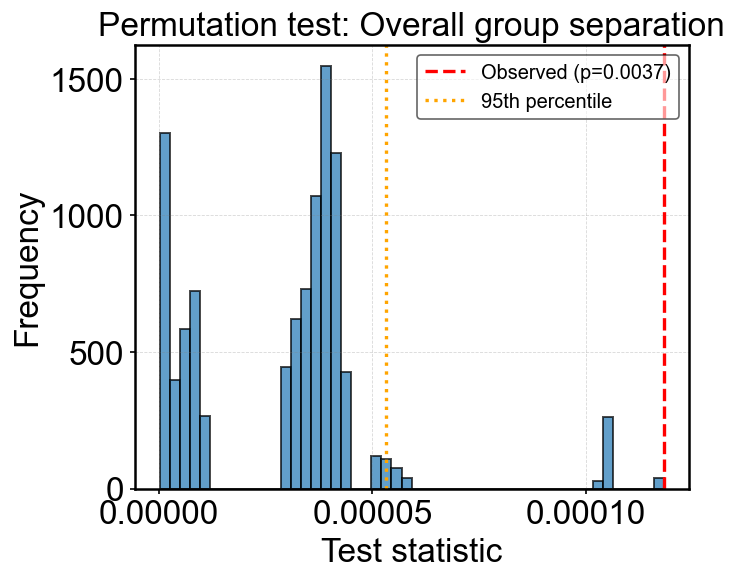

**Supplementary Figure 1**: **Permutation test for overall PCA group separation of unmethylated, 100% 5-mC, and 100% 5-hmC DNA.** The histogram shows the empirical null distribution of the between-group variance statistic $T$ obtained from $B=10,000$random permutations of group labels across all sample replicates in PCA latent space. The red dashed line indicates the observed statistic $T_{\text{obs}}$computed on the true group assignments. The orange dotted line marks the 95th percentile of the null distribution. The observed statistic falls far beyond the null distribution, yielding $p=0.0037$, indicating that the separation of the three groups in PCA space is highly unlikely to have arisen by chance.

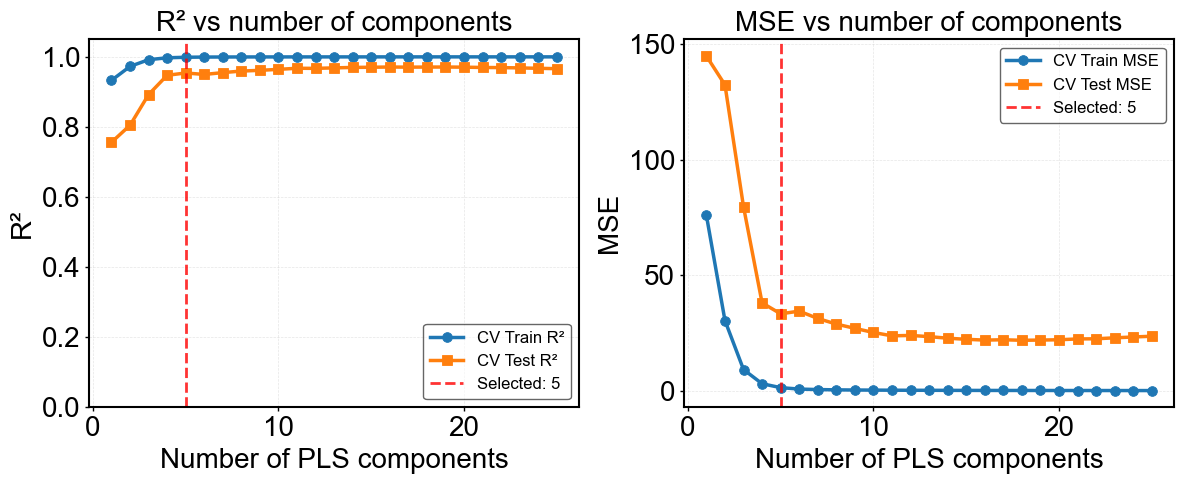

**Supplementary Figure 2**: Optimisation of the number of latent variables (LVs) for PLSR models of individually changing methylation percentage DNA samples, evaluated using the coefficient of determination (R^2^) and mean squared error (MSE). The optimal LV number was determined by cross-validation on the CV train/test split, and model significance was assessed using Van der Voet’s F-test.

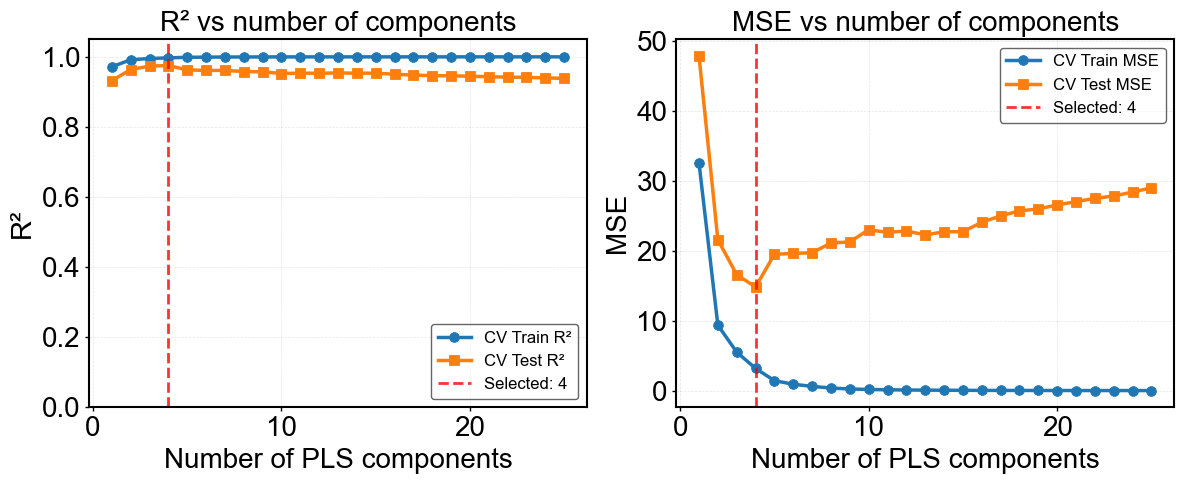

**Supplementary Figure 3:** Optimisation of the number of LVs for PLSR models of individually changing hydroxymethylation percentage DNA samples, evaluated using R^2^ and MSE.

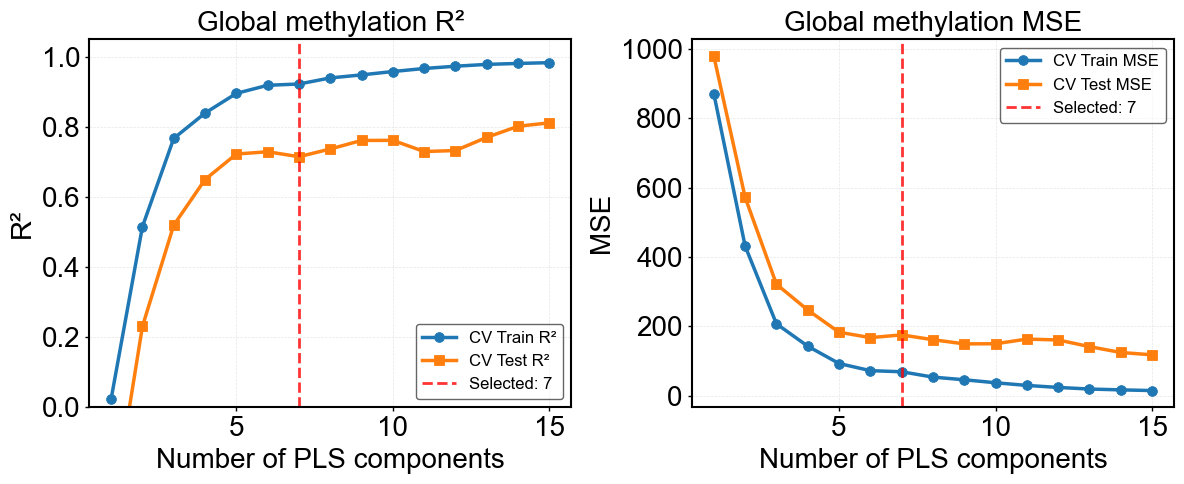

**Supplementary Figure 4:** Optimisation of the number of LVs for PLSR models of globally changing methylation percentage DNA samples, evaluated using R^2^ and MSE.

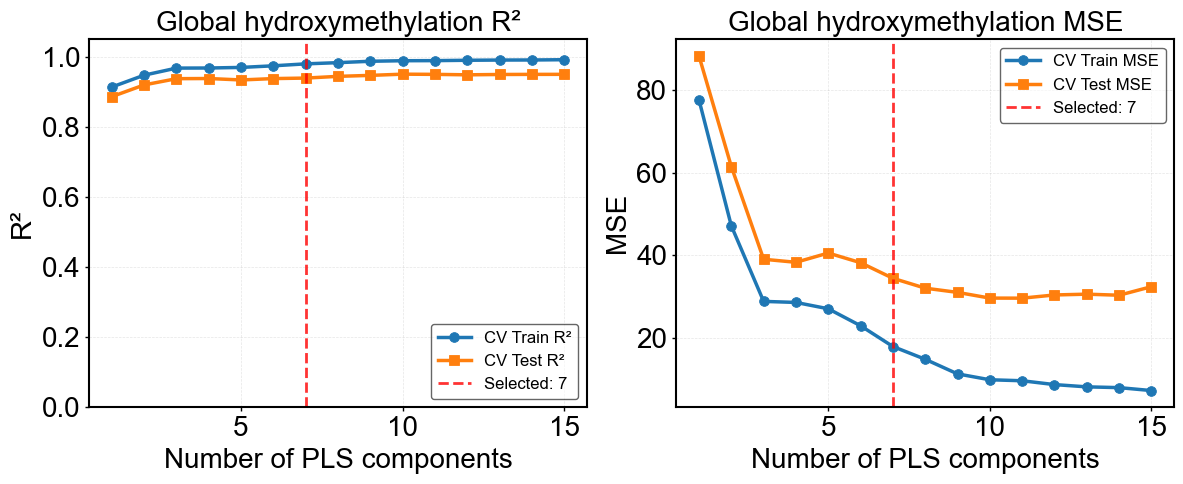

**Supplementary Figure 5:** Optimisation of the number of LVs for PLSR models of globally changing hydroxymethylation percentage DNA samples, evaluated using R^2^ and MSE.

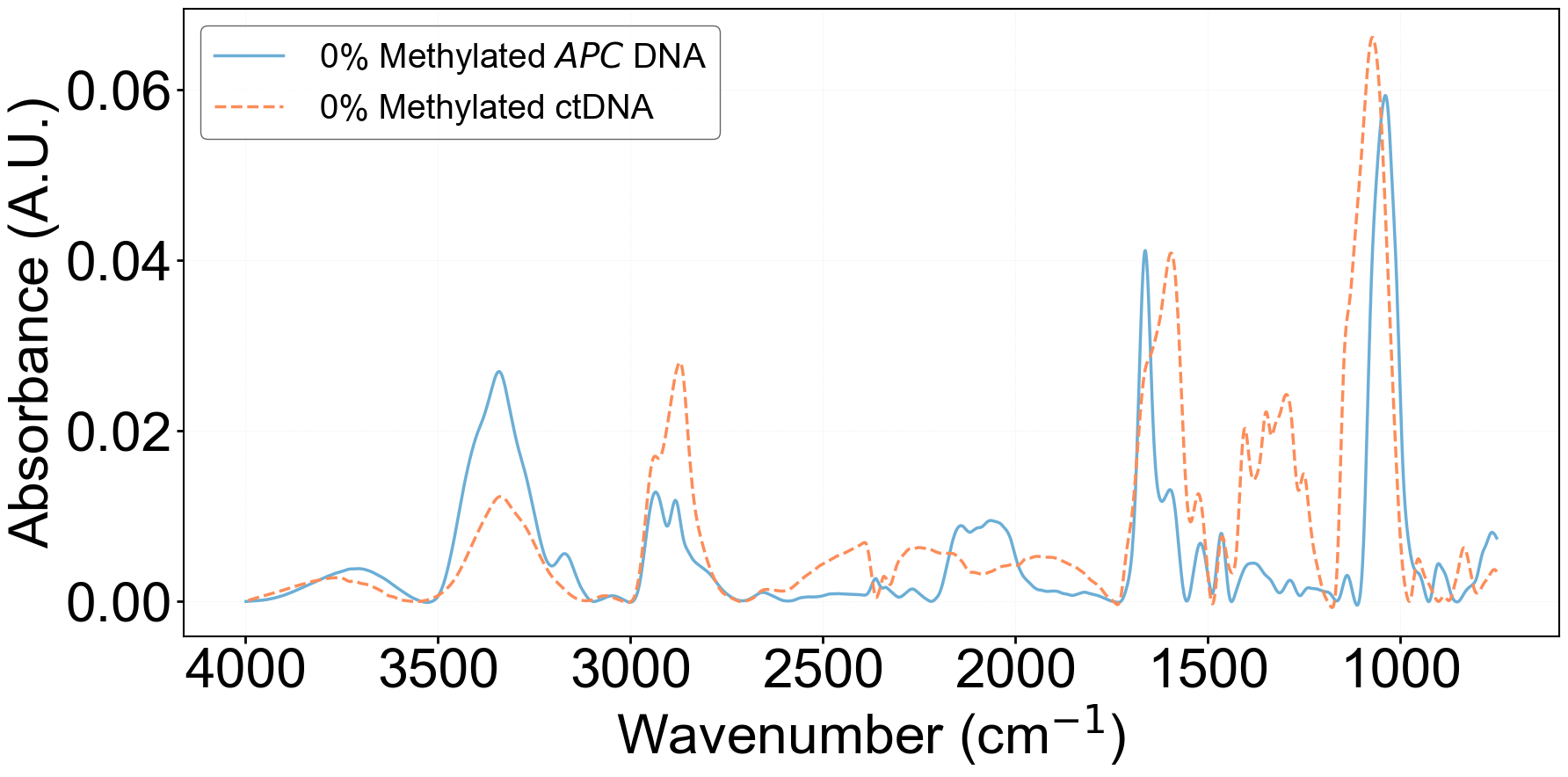

**Supplementary Figure 6:** Comparison of ATR-FTIR spectra of unmethylated *APC* DNA and the unmethylated ctDNA reference standard.
